# Off-the-shelf image analysis models outperform human visual assessment in identifying genes controlling seed color variation in sorghum

**DOI:** 10.1101/2024.07.22.604683

**Authors:** Nikee Shrestha, Harshita Mangal, J. Vladimir Torres-Rodriguez, Michael C. Tross, Lina Lopez-Corona, Kyle Linders, Guangchao Sun, Ravi V. Mural, James C. Schnable

## Abstract

Seed color is a complex phenotype linked to both the impact of grains on human health and consumer acceptance of new crop varieties. Today seed color is often quantified via either qualitative human assessment or biochemical assays for specific colored metabolites. Imaging-based approaches have the potential to be more quantitative than human scoring while lower cost than biochemical assays. We assessed the feasibility of employing image analysis tools trained on rice (*Oryza sativa*) or wheat (*Triticum aestivum*) seeds to quantify seed color in sorghum (*Sorghum bicolor* ) using a dataset of > 1,500 images. Quantitative measurements of seed color from images were substantially more consistent across biological replicates than human assessment. Genome-wide association studies conducted using color phenotypes for 682 sorghum genotypes identified more signals near known seed color genes in sorghum with stronger support than manually scored seed color for the same experiment. Previously unreported genomic intervals linked to variation in seed color in our study co-localized with a gene encoding an enzyme in the biosynthetic pathway leading to anthocyanins, tannins, and phlobaphenes – colored metabolites in sorghum seeds – and with the sorghum ortholog of a transcription factor shown to regulate several enzymes in the same pathway in rice. The cross-species transferability of image analysis tools, without the retraining, may aid efforts to develop higher value and health-promoting crop varieties in sorghum and other specialty and orphan grain crops.

## Introduction

Sorghum (*Sorghum bicolor*) is a grain crop originally domesticated in Africa and now grown across the globe (Fuller and Stevens 2018; Morris et al. 2013a). Sorghum plays a key role in meeting the dietary needs of 500 million people, primarily in Africa, South Asia, and the Americas (Srinivasa Rao et al. 2014). Cultivated sorghum retains high levels of genetic and phenotypic diversity (Mace et al. 2013; Boyles et al. 2019; Boatwright et al. 2022) including substantial variation in grain color (Supplemental Figures S1, S2). Variation in sorghum grain color can indicate variation in the identity and abundance of multiple specialized metabolites present in grain (Wu et al. 2019; Yang et al. 2022).

The colored metabolites present in sorghum grain include anthocyanins, tannins, carotenoids, and phlobaphenes (Davis et al. 2019). Condensed tannins, proanthocyanidins, are antioxidant brown pigments derived from flavan-3-ols (Dixon et al. 2005) and have been linked to multiple desirable human health outcomes, reduced loss of seed to birds (Xie et al. 2019). However, tannins in sorghum are also linked to reduced feed efficiency in livestock and astringent flavors of variable desirability for human food applications. A large proportion of the variation in tannin accumulation in sorghum seeds is explained by two cloned and characterized sorghum genes *tan1* and *tan2* with duplicate recessive interaction controlling the presence of tannins in a layer of cells within the sorghum seed layer called the testa (Wu et al. 2012, 2019). A third uncloned and unmapped gene *spreader* determines whether tannins diffuse into the pericarp (Blakely et al. 1979). In the absence of the *spreader* gene, sorghum seeds with high concentrations of tannin can appear white, yellow, or red, rather than brown. Whether a given sorghum variety will produce white, yellow, or red seeds is determined, at least in part, by two loci, *Y* and *R* (Kambal and Bate-Smith 1976; Zanta et al. 1994). The *Y* locus has been mapped to a gene, *yellowseed1* which encodes a MYB transcription factor homologous to *pericarp color1* (*p1*) in maize (*Zea mays*) (Chopra et al. 1999). Sorghum plants carrying a functional copy of *Y* accumulated significant quantities of the flavan-4-ol derived pigments luteolinidin (orange) and apigeninidin (yellow) (Boddu et al. 2005; Ibraheem et al. 2015). At least three closely related genes or pseudogenes encoding potentially complementary transcription factors are present at the *Y* locus (Nida et al. 2019). Sorghum carrying a dominant haplotype of *Y* and homozygous for recessive alleles at the *R* locus can produce yellow seeds, frequently converting to a tan or brown appearance with age (Dykes et al. 2009). In the absence of other pigment molecules, sorghum varieties homozygous for the recessive haplotype of *Y* will typically produce white seeds. Sorghum varieties carrying dominant alleles of both *Y* and *R* produce red phlobaphene pigments from the flavan-4-ols luteoforol and apiferol, the precursors of luteolinidin and apigeninidin respectively (Chopra et al. 1999; Ibraheem et al. 2015). Several reports suggest *R* may be located on the long arm of chromosome 3 (Mace and Jordan 2010). Yellow seed color in sorghum can also result from the accumulation of yellow carotenoid pigments (Fernandez et al. 2008) regulated by variation in several loci, likely including the phytoene synthase encoding gene *psy3* (Fernandez et al. 2008; Cruet-Burgos and Rhodes 2023). Several genes without direct roles in metabolism are also known to alter apparent seed color in sorghum. These include the action of the *spreader* locus controlling the visibility but not the presence of tannin in sorghum seeds and *Z*, associated with variation in the thickness of the mesocarp, the middle layer of the pericarp, resulting in visible seed phenotypes (Mace and Jordan 2010).

Genetic investigation of the basis of variation in sorghum seed color largely employs data generated via one or more of three approaches: human visual assessment, biochemical quantification, and computer vision. Color data from visual classification was sufficient to identify broad genomic intervals corresponding to *tan1* and *Y* (Morris et al. 2013b) using data from the sorghum association panel, a widely used diversity panel typically consisting of 350-400 sorghum genotypes (Casa et al. 2008). Direct quantification of condensed tannins in the same sorghum association panel was able to localize the position of *tan1* more precisely (Rhodes et al. 2014). An analysis conducted with a much larger population of 1,386 sorghum genotypes and human visual assessment of seed color identified two signals, one corresponding to the *Y* locus and the other which may correspond to the *Z* locus as the same genomic interval was associated with variation in both seed color and mesocarp thickness (Hu et al. 2019). In a smaller population of approximately 250 Chinese sorghum genotypes where high-density marker data was available from whole genome resequencing, a combination of visual assessment of seed color phenotypes and biochemical quantification of tannin concentration was sufficient to identify *Y* and *tan1* (Zhang et al. 2023). Biochemical characterization of the abundance of multiple carotenoids in the seed of the lines of the sorghum association panel identified several signals including one corresponding to zeaxanthin epoxidase (Cruet-Burgos et al. 2023). Measurements of seed color made by extracting the RGB values of pixels corresponding to five seeds per genotype in photos of seeds from the sorghum association panel also identified signals from *Y* and *tan1* (Zhang et al. 2015). A more automated computer vision-based approach using the GRABSEEDS software package implemented within JCVI (Tang et al. 2024) was also able to identify several QTL peaks, including peaks corresponding to *tan1* and *Y*, when employed to quantify the seed colors of a set of several hundred *BC*_1_*F*_2_ families (Nabukalu et al. 2021).

Computer vision-based phenotyping of seed color has the potential to be more quantitative than human visual assessments while being lower cost and higher throughput than biochemical characterization-based methods. However, many manual or semi-automated approaches to quantifying seed color from images are labor-intensive at the image acquisition and/or image annotation stage. In addition, it is unclear how the accuracy and utility of human visual assessment color data, which tends to be qualitative in nature, compares with computer vision-based measures of color which have the potential to be quantitative in nature. Here we assess the feasibility of using pre-trained published seed phenotyping models from other grain crops (Toda et al. 2020) in sorghum without retraining. These models make it possible to identify seeds and extract seed phenotype data from scans generated by spreading sorghum seeds on a flatbed scanner without any need for either ensuring seeds do not touch or manual annotation after image acquisition. We utilize the high throughput quantitative assessments to phenotype seed color across many seeds per entry and conduct genome-wide association studies on a large sorghum diversity panel, to demonstrate that quantitative assessments of seed color from computer vision-based approaches recover more and more strongly supported genetic loci than do qualitative assessments of seed color from human visual assessment. In addition to signals likely corresponding to *y1*, *tan1* and *tan2*, we identify several additional sorghum genomic intervals strongly associated with variation in seed color.

### Core Ideas

- Pre-trained computer vision models can transfer across grass species without retraining.
- Seed color phenotypes quantified from images were more consistent than human assessments of color.
- GWAS conducted using seed color phenotypes from images outperformed GWAS conducted human-scored colors.
- We identified multiple previously unreported GWAS signals near plausible candidate genes for seed color.

## Materials and methods

### Plant Material and Experimental Design

A set of 915 sorghum genotypes drawn from the Sorghum Diversity Panel (Griebel et al. 2021) and Sorghum Association Panel (Casa et al. 2008) were grown at the University of Nebraska-Lincoln’s Havelock Farm in the summer of 2021 in a field with corn planted the previous year. The field employed was located at N*^◦^*40.858, W*^◦^*96.596. The experiment was laid out in a randomized complete block design with two blocks of 966 plots each resulting in a total of 1,932 plots. Each block included one entry of each genotype as well extra replicates of the line BTx623 and Tx430 as repeated checks. Each plot within the field consisted of a single 5-foot (1.5 meters) row with 30-inch (0.76 meters) spacing between parallel rows and 30-inch (0.76 meters) spacing between sequential ranges. Within rows, sorghum seeds were planted at a target spacing of 3 inches (7.62 centimeters) for a target plant density of 21 sorghum plants per row. Before planting, the field received nitrogen fertilizer with a target application rate of 80 pounds of nitrogen per acre (approximately 89 kilograms/hectare) and a pre-emergent application of the herbicide atrazine within 24 hours of planting. Planting occurred on May 25, 2021.

### Seed Image Acquisition and Preprocessing

All grain-bearing panicles from two plants per plot were harvested on October 18th, 2021. Edge plants were avoided when possible. Thirty-four plots did not flower and for an additional 92 plots, a flowering date was recorded during the growing season but no mature seeds were collected. Seeds were removed from panicles using a mechanical thresher and cleaned of chaff and other debris. A qualitative assessment of the color of sorghum seed was recorded. The individuals recording qualitative seed colors were provided with a set of representative seeds of sorghum representing eight color classes (white, grey, yellow, mustard, orange, red, brown, and black) as a standardized reference (Supplemental Figure S2).

For each of the 1,647 plots, a variable number of seeds were loaded onto a flatbed scanner (Epson Perfection V600 Photo with black background) and imaged at a resolution of 300 pixels per inch/dots per inch. Labels with plot and genotype information were included in the area of each scan to reduce the risk of errors. After scanning, a portion of each scan with dimensions of 1,170 *×* 1,150 pixels (9.9 centimeters *×* 9.7 centimeters), which contained sorghum seeds but excluding the scanned label, was extracted.

Sorghum seed scans were preprocessed as previously described (Toda et al. 2020) implementing a pipeline using the OpenCV library (Bradski 2000). A Gaussian blur with an empirically derived 5 *×* 5 window size was applied to each scan to reduce background noise. Blurred color images were converted to eight-bit (0-255) grayscale images using a weighted average value of all three color channels (default R channel weight: 0.299, G channel weight: 0.587, B channel weight: 0.114) in COLOR_BGR2GRAY function in OpenCV. An empirically derived threshold of 45 was used to generate binary image masks from the blurred grayscale images. The OpenCV bitwise_AND operation between the Gaussian blurred color images and binary images was employed to create the final images for downstream analysis. Cropped, blurred, and thresholded images were segmented using two previously published instant ce segmentation neural networks trained on rice and wheat seeds (Toda et al. 2020). Inference of sorghum seed scans via pre-trained models was performed with Tensorflow v.2.10.0 (Abadi et al. 2015) and python using codes available in https://github.com/NikeeShrestha/SorghumSeedSegmentation.

### Model Evaluation and Comparison

Model performance was assessed by comparison to manual segmentation results for approximately 1,600 sorghum seeds across ten images annotated using the Make Sense online annotation platform (Skalski 2019). Recall, (true positives/(true positives + false negatives) was calculated on a per image basis using the compute_recall function provided by Mask_RCN (He et al. 2017) with a threshold of 0.5 intersection/union. An additional set of seeds was drawn from research stocks for 30 sorghum genotypes scanned and processed as described above, and hundred seed weight was determined by counting and weighing 100 seeds to enable comparisons of scanner-derived seed area measurements and conventional measurements of kernel mass.

### Quantification of Sorghum Seed phenotypes

After segmentation via pre-trained models, three seed shape phenotypes: length, width, and area, were quantified using the *skimage*.*measure*.*regionprops*_*table* function implemented in the scikit-image package (Van der Walt et al. 2014) and three color phenotypes: red, green, and blue intensity were quantified using a custom Python script. Seed area was defined as the number of pixels that belong to a given seed instance mask; seed length was defined as the longest distance between pixels included in the seed instance mask along the major axis, and the width was defined as the longest distance between pixels included in the seed instance mask along the minor axis. The red intensity was defined as the average value (between 0 and 255) for the red channel across all pixels included within an individual seed mask. Green and blue intensities were defined equivalently for the respective color channels.

For each of these six phenotypes (seed length, seed width, seed area, red intensity, green intensity, and blue intensity), the average individual values for all seeds in a given image were calculated and assigned to the corresponding sorghum plot. Data from 23 plots were dropped when manual examination of extreme values identified aggregation of many seeds into a single large mask. Scans for 21 plots were removed when visual examinations prompted by large reported differences in seed color between multiple scans of the same sorghum genotype determined the seeds scanned could not have come from the same genotype. A total of 1,603 sorghum seed scans remained after all image-level quality control steps which included one or more seed scans from 881 unique genotypes (Supplemental Data Set S1). After image-level quality control, three principal components (PC1, PC2, and PC3) of variation for seed color phenotypes were calculated from the average red, green, and blue intensity phenotypes described above using the built-in *princomp* function in R v.4.2.0 (R Core Team 2020). Visual examination of phenotype distributions on a per-plot basis was used to identify and remove extreme values for each phenotype (Supplemental Figure S3A). This step resulted in the removal of seed length data for one plot, average blue intensity for 11 plots, average green intensity for 7 plots, and PC1, PC2, and PC3 scores for 9, 20, and 12 plots respectively.

### Genetic Marker Information

Genetic markers used in this study were generated using RNA-seq data from mature leaf tissue of plants grown in the same 2021 field experiment employed for phenotyping. RNA sequencing libraries were sequenced on an Illumina NovaSeq6000 with a target read depth of 20 million total sequenced fragments and 2 x 150 base pairs of sequencing per fragment. Raw reads were trimmed using Trimmomatic v0.33 with the following parameters ILLUMINACLIP:TruSeq3-PE-2.fa:2:30:10 LEADING:3 TRAILING:3 SLIDINGWINDOW:4:15 MINLEN:35 (Bolger et al. 2014). Trimmed reads were then mapped to the sorghum BTx623 reference genome V5 (Institute 2023) using STAR v2.7.9a (Dobin et al. 2013) with parameter settings of outFilterMismatchNmax 30, outFilterScoreMinOverLread 0.1, outFilterMatchNminOverLread 0.1 and seedSearchStartLmax 20. Initial SNPs were called using Haplotype Caller in GATK4 v4.1 with the following parameters; "QD < 2.0", "QUAL < 30.0", "SOR > 3.0", "FS > 60.0", "MQ < 40.0", "MQRankSum < -12.5" and "ReadPosRankSum < -8.0" (Poplin et al. 2017). These initial SNP markers were filtered to retain SNPs with a minor allele frequency >0.01 and frequency of heterozygous genotypes call <0.1 using VCFtools v.0.1 (Danecek et al. 2011) and bcftools v.1.17 (Danecek et al. 2021). Missing genotype calls in the SNP set were imputed using Beagle v5.2 (Browning et al. 2018). For GWAS, the imputed SNP set was subsampled to retain only SNPs with a minor allele frequency of *>* 0.05 and frequency of heterozygous genotypes calls *<* 0.05 among the common 682 genotypes between population phenotyped and genotyped in the study. These criteria resulted in a set of 169,600 SNPs being retained.

### Quantitative Genetic Analyses

Repeatability for individual seed phenotypes was estimated using phenotype data for 629 genotypes for which phenotype data was collected from both replicated blocks of this study. Repeatability was calculated using the equation:

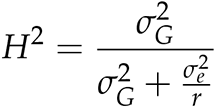

Where 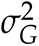 is the total amount of variance explained by genotype and 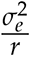 is the total residual variance divided by the number of replications of each genotype. A mixed linear model of the form *y_i_* = *µ* + *Genotype_i_* + *error_i_* was used to estimate the total variance explained by genotype and the total residual variance for each phenotype, where, *y_i_*is the mean phenotype of interest in the genotype, *µ* is the overall mean of the population, *Genotype_i_* is random effect of genotype *i*, and *error_i_* is the residual error. The model was implemented using the lme4 package (Bates et al. 2015) in R v.4.2.0 (R Core Team 2020).

Genome-wide associations were conducted using phenotype average values for 682 sorghum genotypes which were scored in at least one of the two replicated blocks and for which genetic marker data was also available. Manual examination of the distributions of genotype-level averages (Supplemental Figure S3B) led to the removal of genotype level data for seed area (3 genotypes), seed width (7 genotypes), average red intensity (29 genotypes), average green intensity (20 genotypes), average blue intensity (7 genotypes), and data for 5, 17, and 7 genotypes for PC1, PC2, and PC3 of color intensity respectively.

GWAS was conducted using the FarmCPU GWAS algorithm implemented in the rMVP software package (Liu et al. 2016; Yin et al. 2021). The first three principal components of variance calculated from the genetic marker data were included as covariates, and the genomic relationship matrix was included to account for population structure. This GWAS analysis was implemented with resample model inclusion probability (RMIP) (Valdar et al. 2009). One hundred iterations of FarmCPU GWAS analysis were conducted for each phenotype and in each iteration, a random 10% of sorghum genotypes were masked, and a separate FarmCPU GWAS analysis was conducted. In each iteration, the threshold for an SNP to be considered significantly associated was *p–value* less than 9.70*×*10*^−^*^7^. This threshold was calculated using a Bonferroni corrected 0.05 p-value cutoff considering the estimate of 51,509 independent genetic marker numbers in the genetic marker dataset employed in this study obtained from GEC v.0.2 (Li et al. 2012). SNPs identified in at least 10 of the 100 resampling GWAS analyses (RMIP *≥* 0.1) were considered significant associations in the final analysis.

## Results

Two models pre-trained on seeds from other grain crops (rice or wheat) (Toda et al. 2020)) both successfully identified the majority of sorghum seeds present in ten manually annotated images generated by scanning seed samples from ten different sorghum genotypes (Figure 1A). While the performance of both models was imperfect, the performance issues presented by the two models were different. The model trained on rice seeds generated masks that often excluded portions of individual seeds visible in the image (Figure1B). The model trained on wheat seeds generated more complete masks for seeds it identified, but more frequently failed to identify a significant percentage of the sorghum seeds present in images (Figure1C). At the single seed level, the area of seeds as estimated from automated masks was highly correlated with the area of seeds as estimated from manually annotated seed masks (rice model: *R*^2^=0.93, wheat model *R*^2^=0.94) (Figure 1D&E).

**Figure 1.**
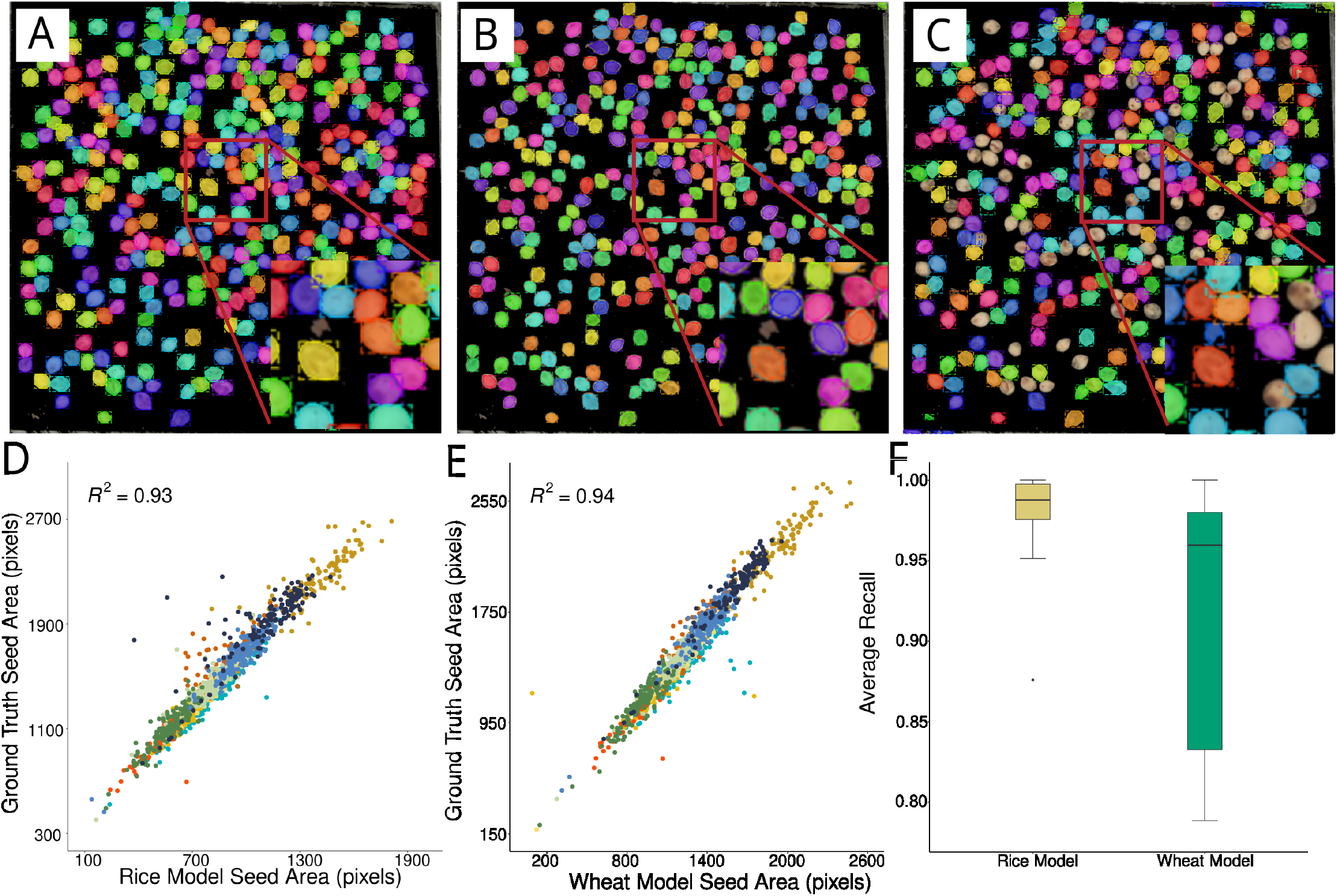
Comparison of the performance of models trained on rice and wheat seeds at the task of identifying and segmenting sorghum seeds. A) Example of the manually annotated seed positions used as ground truth in this study. B) Example of seed positions and shapes identified by the model trained on rice seeds. C) Example of seed positions and shapes identified by the model trained on wheat seeds. D) Relationship between manually annotated seed area and automated seed area measurements obtained from the rice-trained model. Each point indicates a single seed which was identified via both manual and automated annotation. Different colors represent different seed scans. E) Relationship between manually annotated seed area and automated seed area measurements obtained from the wheat-trained model. Different colors represent different seed scans. F) The average recall with 0.5 Intersection/Union in each of the ten images which were identified via masks generated using either the pre-trained rice model or the pre-trained wheat model.

The issue with the model trained on wheat seeds failing to identify some sorghum seeds appeared to be either image or genotype-specific with >94% of manually annotated sorghum seeds correctly identified in 7/10 images, but only 80% of manually annotated sorghum seeds correctly identified in the remaining three images (Figure 1F). Seed area estimated from both models exhibited an equal correlation (*R*^2^) of 0.77 with ground truth measurements of 100-grain mass for a separate set of 30 sorghum genotypes selected to represent the full range of seed sizes observed in the sorghum diversity population (Supplemental Figure S4A, B). Given the consistent performance of the two models in estimating variation in seed area and the relatively poorer performance of the wheat-trained model in estimating total seed counts and low average recall, the outputs of the rice-trained model were employed for subsequent analyses.

Three seed-shape phenotypes (seed area, length, and width), three seed color phenotypes (average red, green, and blue intensities), and three principal components of variation in seed color (PCs) were extracted from each image using the masks generated using the model trained on rice seeds. Substantial variation was observed for each seed shape and color phenotype as well as for the three principal components derived from three color channels (Supplemental Table S1). The first principal component explained the highest proportion of total variance, 96.74%, the second principal component explained 2.78% of the total variance and the third principal component explained 0.46% of the total variance. The genetic repeatability (*H*^2^) for six seed-related phenotypes extracted after seed segmentation ranged from 0.91 to 0.94 with average blue intensity having the highest *H*^2^ of 0.94 and seed area and length having the lowest *H*^2^ of 0.91.

The seeds assigned to the qualitative color categories brown, orange, yellow, gray, and white by human scorers exhibited different but overlapping distributions across the first two principal components of variation in seed color (Figure 2). A number of individual sorghum seed samples whose manual categorical color classification and PC phenotype values were inconsistent with each other were manually rechecked and visual inspection of scans found seed colors that were consistent with PC scores rather than manual color category assignment (Supplemental Figure S5). Discordance between manual and imaging-based seed phenotyping tended to occur in samples with either staining or mold on the seed surface or with scattered brown/red dots across the seeds (Supplemental Figure S5). Notably, when asked to classify sorghum seeds into one of eight color categories, replicates of the same sorghum genotype grown in different parts of the field were assigned different color categories 40% of the time, although this declined to 8.5% when the manual color annotation was reduced to a two category system; light (including the original categories "white", "grey", "mustard", and "yellow") and dark ("orange", "red", "brown" and "black") system (Figure 3).

**Figure 2.**
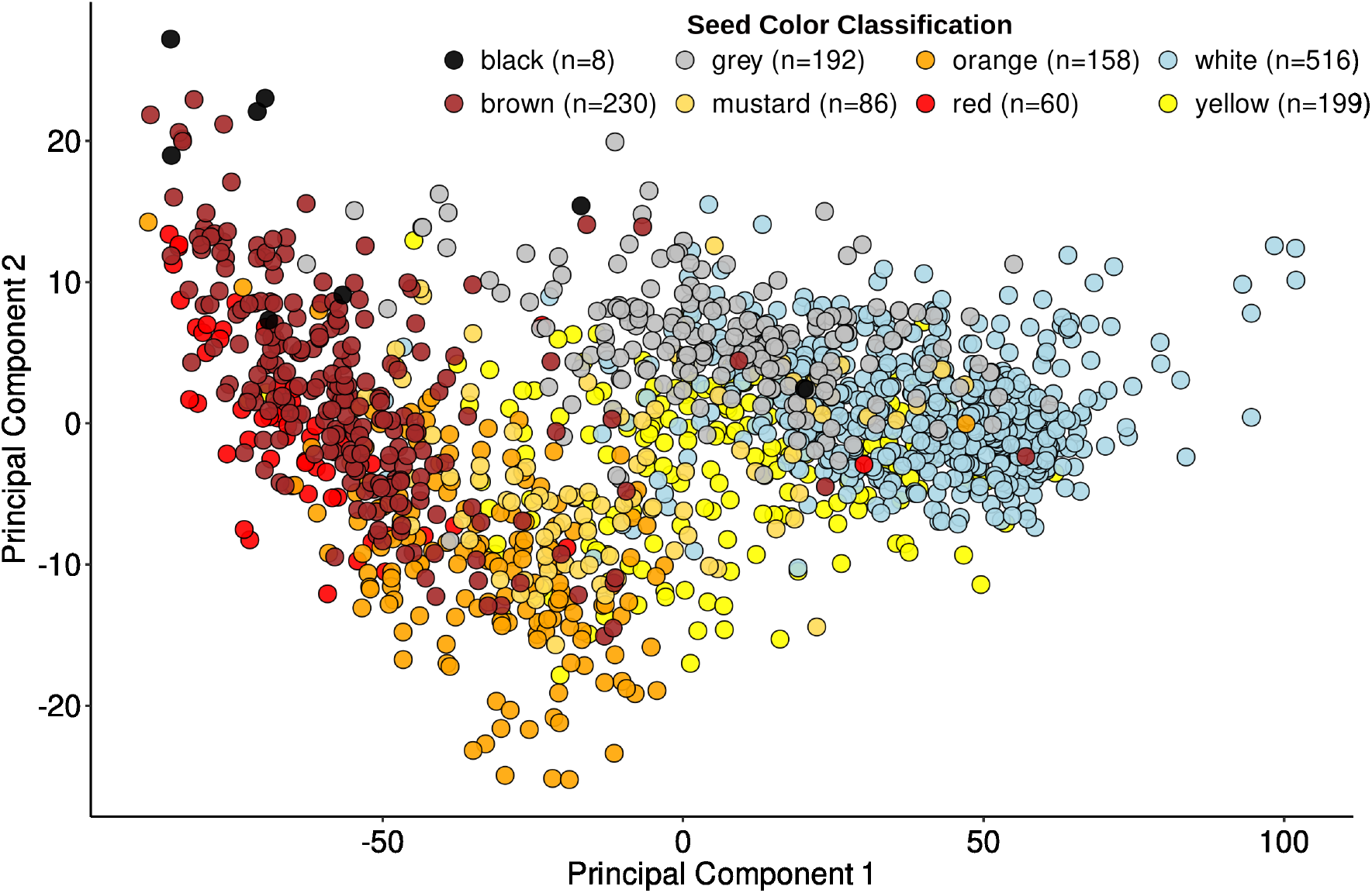
Relationship between qualitative ground truth color classification and quantitative measurements of sorghum seed color. Distribution of scores for the first two principal components of variation in color phenotypes (average red, blue, and green intensity) for scans of 1,449 sorghum plots grown and harvested as part of this experiment. The colors of individual points indicate the categorical color phenotype assigned to each plot during manual seed color phenotyping.

**Figure 3.**
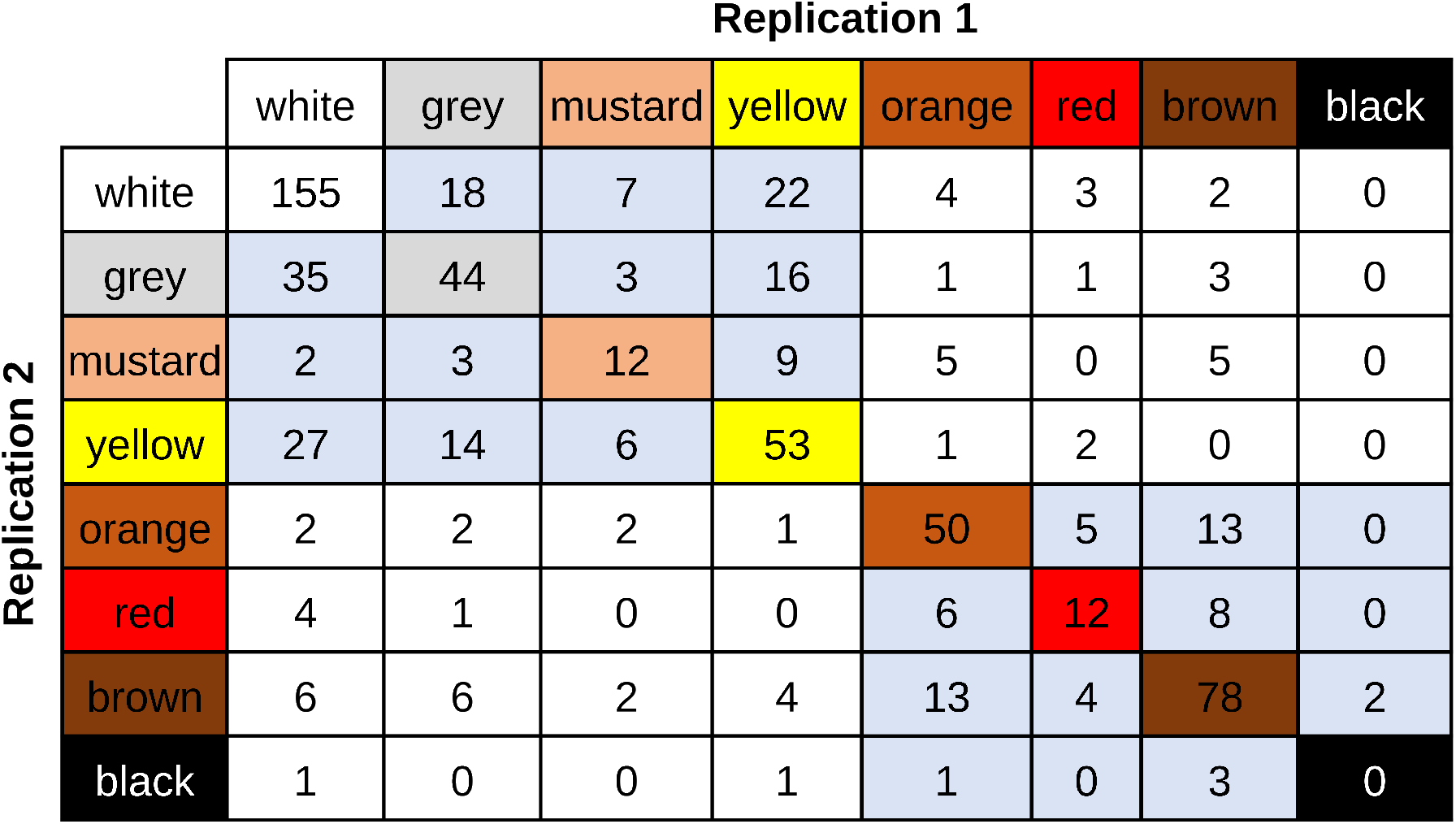
Disagreements in color classification between independent replicates of the same sorghum lines grown in the same field experiments. Data shown are for 680 sorghum genotypes for which qualitative color scores were recorded in from two independently replicated plots in 2021. Numbers in colored boxes indicate exact color category matches in the eight-color system (N=404). Numbers in light blue boxes indicate disagreements between replicates in color category assignments in the eight-color system which still match in the two-color category system (light = white, gray, mustard yellow, dark = orange, red, brown, black) (N=217). Numbers in white boxes indicate disagreements between replicates under both the eight-color system and the two-color system (N=59).

We employed a set of genetic markers scored for 682 of the 881 sorghum genotypes phenotyped above to conduct genome-wide association studies for seed shape and color phenotypes. Resampling-based analysis of FarmCPU GWAS identified 19 significant marker-trait associations representing 15 unique markers associated with seed area (7 significant associations), seed length (4 significant associations), and/or seed width (8 significant associations) above a resampling model inclusion probability (RMIP) threshold of *≥* 0.1 (Figure 4, Supplemental Data Set 2A). One of the 15 unique genetic markers was associated with both seed area and length, and one was associated with seed length, seed area, and width. Two of the 15 unique genetic markers identified in GWAS for seed shape phenotypes overlapped and one genetic marker was identified 11 kb downstream of previously reported associations from a sorghum NAM population and/or sorghum association population and on (Tao et al. 2020). Four of the 15 genetic markers significantly associated with sorghum seed size or shape phenotypes were located within 400 kilobases of the sorghum orthologs rice or maize genes previously linked to variation in seed size or shape (Figure 4).

**Figure 4.**
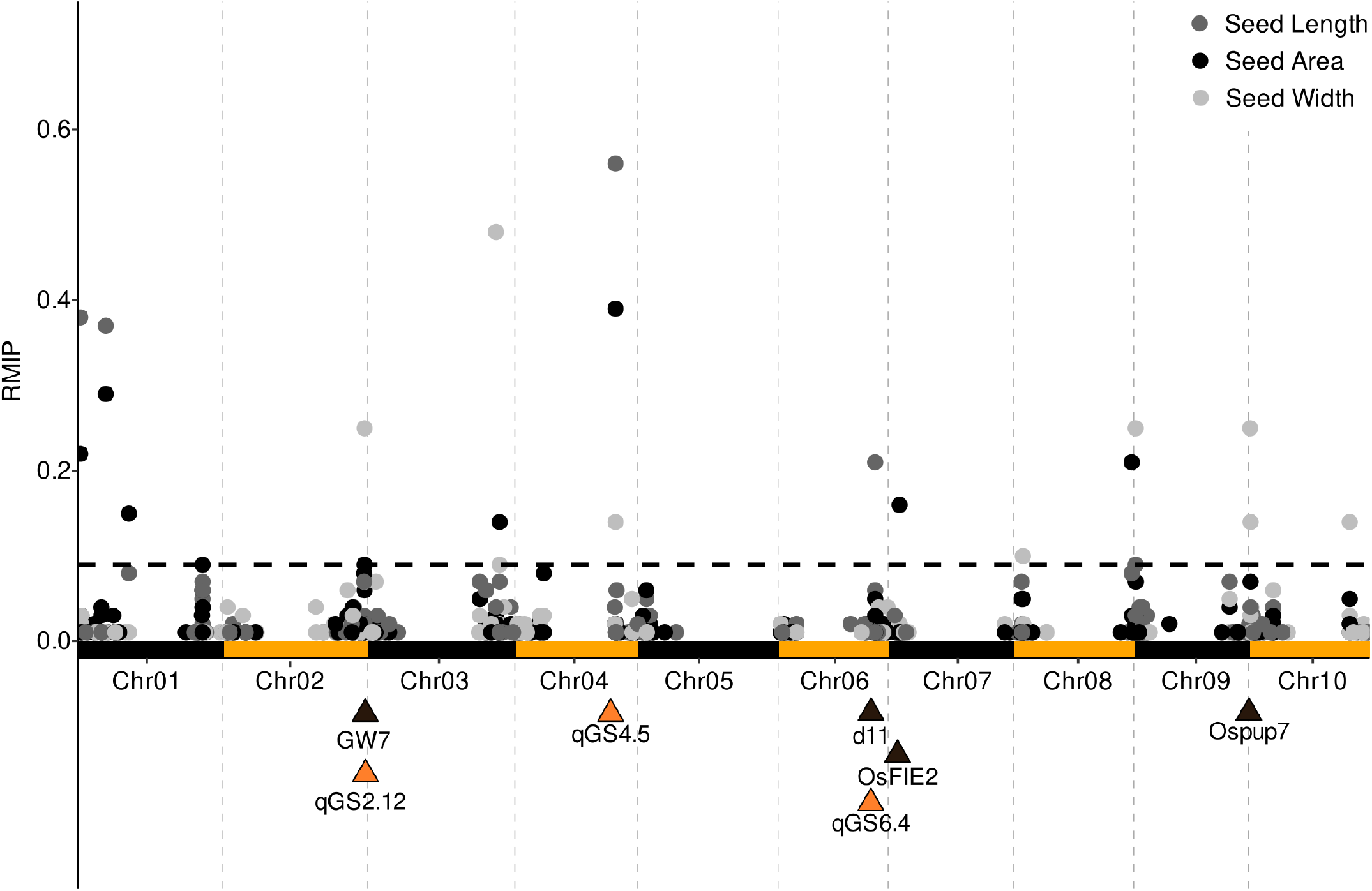
Genetic markers associated with variation in seed shape phenotypes. Results of a resampling-based GWAS analysis conducted for three seed shape phenotypes: seed area, seed length, and seed width. Each point indicates the position of an individual genetic marker (x-axis) and the proportion of 100 FarmCPU GWAS iterations in which that marker was significantly linked to variation in the phenotype of interest (Resampling Model Inclusion Probability, RMIP). The horizontal dashed blast line indicates a threshold of RMIP *≥* 0.1. Yellow triangles indicate the positions of previously described QTL for seed shape or size in the sorghum NAM population and/or sorghum association population (Tao et al. 2020). The QTLs, qGS2.12 and qGS4.5, overlap with the significant genetic markers and qGS6.4 is 11 kb upstream of the genetic marker. Black triangles indicate the positions of sorghum orthologs of rice or maize genes linked to seed shape or size located within 400 kilobases of a marker associated with seed shape phenotypes. Sorghum gene; Sobic.002G367300; ortholog of *GW7* in rice (Wang et al. 2015), Sobic.006G114600; ortholog of *d11* in rice and maize (Sun et al. 2021), Sobic.007G032400; ortholog of *OsFIE2* in rice (Folsom et al. 2014), Sobic.009G227201; ortholog of *Ospup7* in rice (Ji et al. 2019).

A genome-wide association study for manually classified sorghum seed color phenotypes – two color categories – identified a total of ten significant marker-trait associations including two that likely correspond to the known sorghum color genes *y1* (Sobic.001G398100) and *tan1* (Sobic.004G280800) (Figure 5A, Supplemental Data Set 2B). Both genes were associated with the only two markers associated with manually scored color data with RMIP *≥* 0.5. Genome-wide association studies conducted for six different quantitative color phenotypes extracted from scanned and segmented seed images identified a total of 70 marker-trait associations consisting of 43 unique genetic markers significantly associated with one or more quantitative color phenotypes (Figure 5B, Supplemental Data Set 2C). Out of the 11 genetic markers associated with multiple color phenotypes, 6 genetic markers were associated with three color channels and the first principal component, 3 signals were identified to be associated with average blue and green intensities, and the first principal component, one signal was associated with average red and blue intensities and the first principal component and one signal were associated with average red intensity and the first principal component. The signals identified in the analysis using color phenotypes from scanned sorghum seeds included marker-trait associations corresponding to three known color genes, including signals corresponding to the locations of *y1* (Figure 6A) and *tan1* with higher resampling model inclusion probabilities than in the manual color classification based analysis and an additional signal in the general vicinity of *tan2* (Sobic.002G076600).

**Figure 5.**
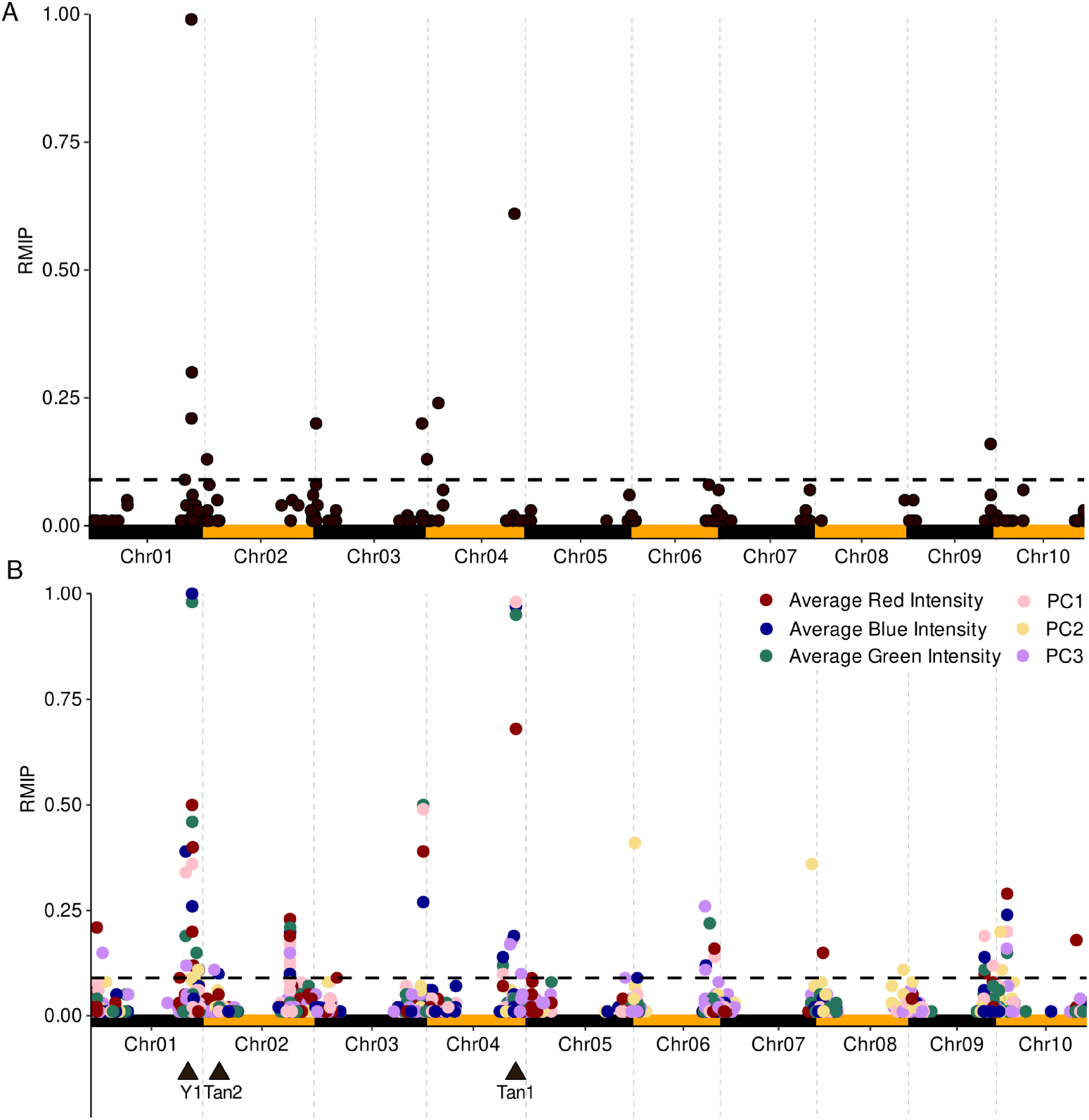
Genetic markers associated with variation in sorghum seed color. A) Results of a resampling-based GWAS analysis conducted by collapsing the eight manually described color cat- egories into two categories as described in Figure 3. **B)** Results of resampling-based GWAS analysis conducted for three quantitative color phenotypes: average red, blue, and green intensity for seed pixels, calculated directly from segmented seed images and the three principal components of variation calculated from those three initial phenotypes. Analysis was conducted using data for 682 sorghum genotypes. Black triangles indicate the positions of previously characterized genes known to play a role in determining variation in seed color in sorghum.

**Figure 6.**
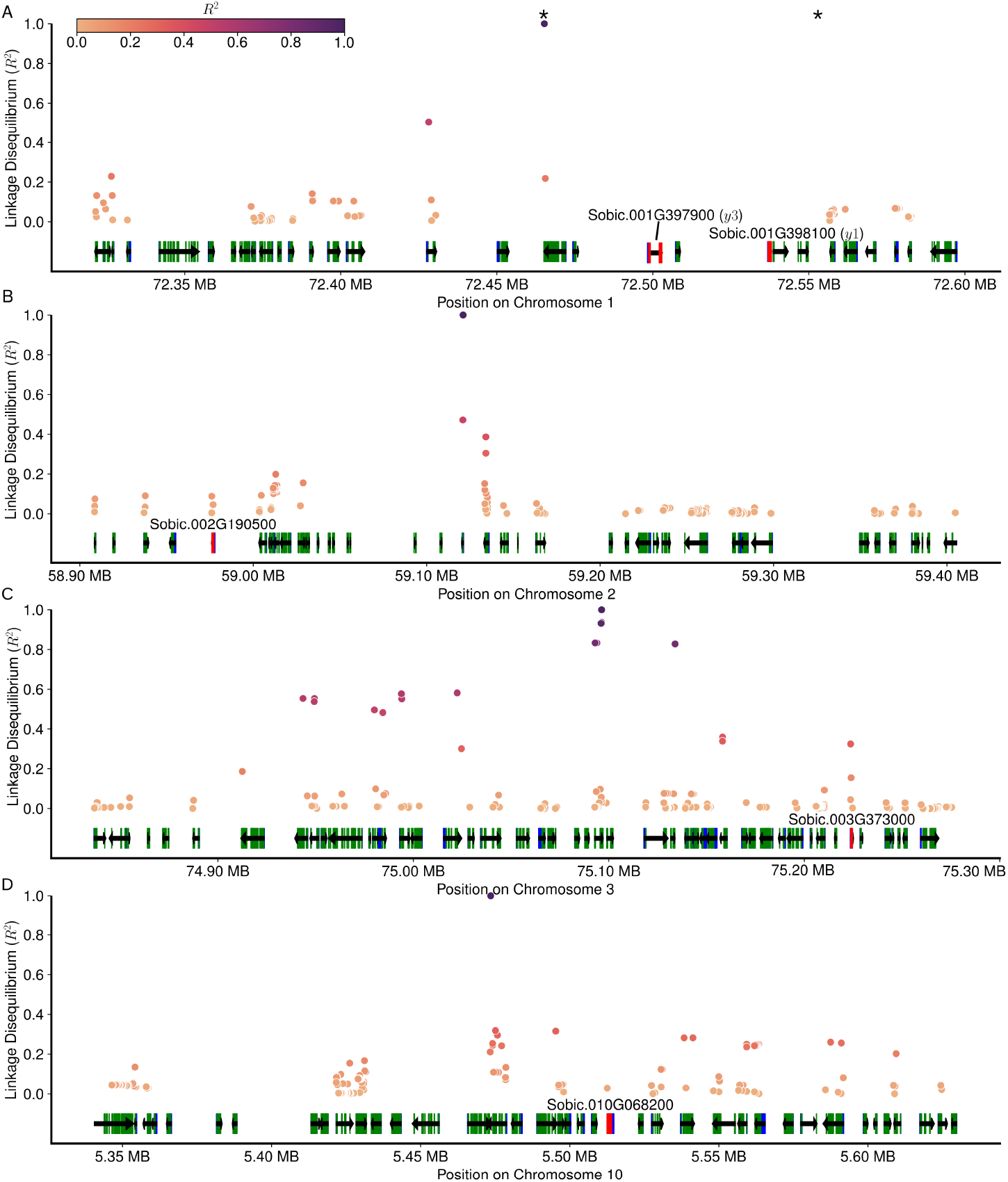
Genomic intervals and phenotypic effects associated with color GWAS hits on chromosomes 1 and 3. A) Linkage disequilibrium and annotated gene models within the genomic interval surrounding the genetic marker (Chr01:72,465,237) associated with variation in both manually scored qualitative sorghum seed color and automatically scored quantitative sorghum seed color. Each point indicates the physical position (x-axis) and linkage disequilibrium with the trait-associated SNP (y-axis) of an SNP within the genomic interval. Black arrows indicate the positions of annotated genes, green boxes the position of protein-coding exons, and blue boxes the positions of untranslated exons. Genes with red exons are *yellowseed1* (*y1*) and *yellowseed3* (*y3*). The positions of two SNPs that were independently associated with seed color (Chr01:72,465,237 and Chr01:72,556,673) are indicated with asterisks. **B)**. Linkage disequilibrium and annotated gene models within the genomic interval surrounding the genetic marker (Chr03:75,096,302). The position of a candidate gene, the ortholog of a rice MYB transcription factor that regulates flavonoid metabolism is marked in red. **C)** Linkage disequilibrium and annotated gene models within the genomic interval surrounding the genetic marker (Chr02:59,121,010). The position of an *α* amylase encoding gene previously reported as a potential candidate gene for the classical *Z* locus and associated with variation in both seed color and mesocarp thickness in sorghum is marked in red. **D)** Linkage disequilibrium and annotated gene models within the genomic interval surrounding the genetic marker (Chr10:5,473,493). The position of a candidate gene, the sorghum ortholog of a rice flavonol synthase/flavanone-3-hydroxylase is marked in red.

Excluding signals in the vicinity of three cloned and characterized color genes *y1*, *tan1* and *tan2*, three additional genetic markers were identified significantly associated with variation in all three color channels as well as the first principal component of variation: Chr02:59,121,0101 (highest RMIP = 0.23), Chr03:75,096,302 (highest RMIP = 0.5), and Chr10:5,473,493 (highest RMIP = 0.29). The signal on chromo- some 2 was in the rough vicinity of a signal previously identified in a GWAS for manually assigned seed color in a much larger sorghum population and 300 kilobases away from Sobic.002G190500 (Figure 6B, Supplemental Data Set S3A), a gene encoding an *α* amylase identified in that study as a potential candidate for the causal gene underlying the *Z* locus (Hu et al. 2019). The signal on chromosome 3 was supported in the categorical human-scored seed color GWAS (RMIP = 0.2) (Figure 5A) in addition to the image-based seed color GWAS. Genetic markers in a genomic interval of 440 kilobases around the chro- mosome 3 hit exhibited moderate to strong linkage disequilibrium (*R*^2^ *≥* 0.25) with the GWAS-tagged marker. This interval contained a total of 38 annotated gene models (Figure 6C, Supplemental Data Set S3B). One of these genes, SbMYB50/Sobic.003G373000, is the ortholog of a MYB transcription factor (LOC_Os01g65370) in rice that has been shown to repress the expression of flavonoid-3-hydroxylase and a chalcone flavonone isomerase (Figure 7) based on evidence from overexpression lines (Grotewold et al. 1994; Sun et al. 2023). The classical, but as yet uncloned, color gene *R* is also believed to be located on chromosome 3 (Mace and Jordan 2010). However, while *Y* is known to be epistatic to R (Kambal and Bate-Smith 1976), the interaction between the effects of the *Y* locus-linked marker and the effects of the chromosome 3 color-linked marker was not significantly different from additive (Supplemental Figure S6C-E). The signal for seed color variation on chromosome 10 is approximately 40 kilobases away from Sobic.010G068200, the sorghum ortholog of a rice gene (LOC_Os10g40880) annotated as either a putative flavonol synthase or flavanone 3-hydroxylase (Figure 6D, Supplemental Data Set S3C). Either of these enzymatic activities would place this gene in the biosynthetic pathway responsible for the synthesis of the majority of known colored metabolites present in sorghum seeds (Figure 7).

**Figure 7.**
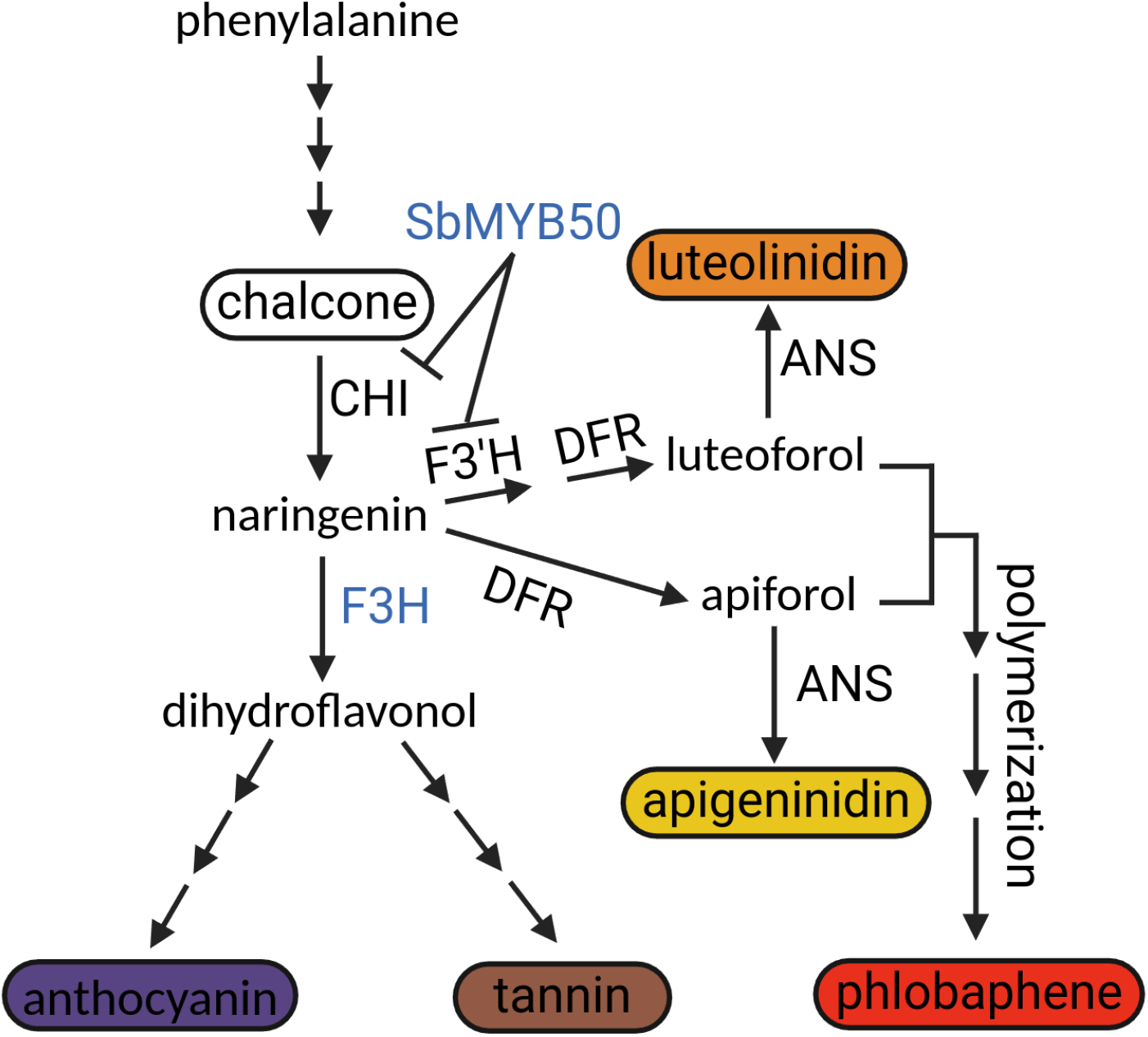
Schematic of a section of phenylpropanoid pathway leading to the synthesis of multiple colored metabolites found in sorghum seeds. Enzyme abbreviations: CHI, chalcone isomerase; F3H, flavanone 3-hydroxylase; DFR, dihydroflavonol 4-reductase; F3H, flavanoid 3-hydroxylase; F3‘H, anthocyanidin synthase; ANS. Multiple arrows represent multiple synthesis steps. Names shown in blue SbMYB50 (Sobic.003G37300), and F3H (Sobic.010G068200) were identified in the vicinity of genetic markers significantly associated with multiple seed color phenotypes. Background colors for pigments indicate known/reported colors from the literature.

## Discussion

When grains are consumed directly, seed color plays a key role in consumer acceptance of new crop varieties. Seed color can also be a marker for bioactive compounds with the potential to improve or impair human health (Yang et al. 2022). However, seed color is frequently still assessed in an *ad hoc* fashion via human classifiers who seek to divide quantitatively varying colors into discrete categories. We found that these human-assigned qualitative color scores had a high rate of discordance (Figure 3, while quantitative color phenotypes extracted from scans of sorghum seeds were highly repeatable (*H*^2^>0.9) across multiple plots of the same sorghum genotypes. This included the identification of marker-trait associations corresponding to three characterized sorghum genes known to influence seed color (*y1*, *tan1*, and *tan2*) (Zanta et al. 1994; Wu et al. 2012, 2019) along with one locus corresponding to the likely location of the classical Z locus, and two other loci near genes with plausible links to anthocyanins, tannins, and/or phlobaphene metabolism (Figure 7).

In GWAS analysis using quantitative seed color phenotypes derived from seed scans, the two marker-trait associations were found in chromosome 1, 91 kb away from each other (Figure 6A). The one association (Chr01:72,465,237, highest RMIP = 1) linked to all three color intensities phenotypes and the first principal component was approximately 71 kb upstream from cloned *y1* gene and another association (Chr01:72,556,673, highest RMIP = 0.46) linked to green and blue color intensity and first principal component was approximately 18 kb downstream of *y1* gene. FarmCPU controls for the effect of previously identified marker-trait associations when evaluating the significance of subsequent markers (Liu et al. 2016) and these two markers also exhibit very low linkage disequilibrium (LD <0.01) suggesting they correspond to different functional variants rather than providing redundant information on the same causal locus. This would be consistent with the previously reported complex architecture of Y locus which has multiple copies of R2R3 MYB genes (*yellowseed3*, *y1* and additional pseudogenes) within the same vicinity with both genes complimentary linked to grain color in sorghum (Nida et al. 2019, 2021). An additional independent signal (Chr01:67,840,021, highest RMIP = 0.39) approximately 4 MB upstream of a signal at Chr01:72,465,237 was identified linked to blue intensity phenotype and the first principal component (Figure 4B) and is approximately 262 kb upstream of previously reported candidate gene (Sobic.001G349900) for variation in exocarp color in Chinese sorghum germplasm (Zhang et al. 2023). Identification of multiple marker-trait associations on chromosome 1 in this and previous studies suggest numerous other loci on chromosome 1 in addition *y1* may contribute to seed color pigmentation in sorghum.

A strong and repeated signal on chromosome 3 was identified in both the analysis of sorghum seed color based on seed scans (highest RMIP = 0.5) and human color assessment (highest RMIP = 0.2). The signal we identified on chromosome 3, is also distinct from a repeatedly reported signal from previous GWAS and QTL mapping studies located at approximately 64 MB (Kimani et al. 2020; Nida et al. 2021; Kumar et al. 2023), 11 MB from the signal we identify on the same chromosome at position 75.09 MB. The large interval ( 400 kb) defined by linkage disequilibrium around this hit includes Sobic.003G373000. Sobic.003G373000 is the ortholog of LOC_Os01g65370, which interacts with TOPLESS and HDAC1 to form a transcriptional repressor complex, which inhibits the expression of two flavonoid-3‘-hydroxylase (F3‘H) and a chalcone flavonone isomerase (CHI) gene in the metabolic pathway leading to production of different pigments in plants (Sun et al. 2023). F3‘H and CHI catalyze reactions at metabolic junctions which can lead to different pigmentation in plants (Grotewold et al. 1994; Falcone Ferreyra et al. 2012). The position of this chromosome 3 GWAS signal and associated candidate gene is somewhat consistent with the reported approximate localization of the uncloned sorghum locus R which acts downstream of *y1* (Mace and Jordan 2010; Rhodes et al. 2014). However, in the absence of strong statistical evidence supporting an epistatic interaction between this marker-trait association on chromosome 3 and *y1* for seed color, as well as inconsistent location of signal with previously reported associations, the location does not yet represent strong evidence for having identified the location of R.

Sorghum produces a wide range of bioactive compounds in grain such as tannins, phenols, antho- cyanins, and carotenoids which are shown to alter the composition of the gut microbiome, including in ways linked to improved outcomes for obesity, diabetes, oxidative stress, cancer, and hyperten- sion (de Morais Cardoso et al. 2017; Yang et al. 2022). Previous efforts with smaller sorghum populations have demonstrated that, in some cases, sorghum loci associated with changes in the abundance of multi- ple beneficial bacterial taxa in the human gut microbiome colocalize with loci associated with variation in seed color (Yang et al. 2022; Korth et al. 2024). Here we have demonstrated that a combination of quantitative measurements of color enabled by computer-vision-based approaches to seed phenotyping with analysis of a substantially larger sorghum population can lead to the identification of new candidate genes SbMYB50 (Sobic.003G373000) and the putative flavonol synthase or flavanone 3-hydroxylase Sobic.010G068200 that may play roles in determining the abundance and identity of bioactive molecules with the potential for beneficial or detrimental impacts on human gut microbiome (Petitot et al. 2017; Korth et al. 2024). These results could serve either as the basis for future efforts to fine map, clone, and characterize the specific genes involved in regulating variation in seed color in sorghum and/or as the basis for marker-assisted selection efforts to develop new sorghum varieties with specific suites of bioactive pigment molecules as a tool to impact human and/or animal health via the human gut microbiome.

This study also tests and validates the potential to deploy pre-trained AI models for image analysis across species within the grasses. Both models trained with rice data or trained with wheat data exhibited acceptable performance on sorghum. This result is consistent with the previous observation that machine learning models trained to semantically segment sorghum plant organs in hyperspectral images also achieved good performance in semantically segmenting maize organs (Miao et al. 2020). Cross-species transferability efforts devoted to developing artificial intelligence models for image analysis in the three-grain crops that provide the majority of the global calorie needs today – rice, wheat, and maize – may also benefit and accelerate crop improvement efforts in many of the other grain crops which currently play smaller roles in the global food supply but exhibit greater resilience and resource use efficiency, including pearl millet (*Cenchrus americanus* syn. *Pennisetum glaucum*) and proso millet (*Panicum miliaceum*) in addition to sorghum (Shrestha et al. 2023; Wimalasiri et al. 2023).

## Supporting information

Supplemental Data Set S1

Supplemental Data Set S2

Supplemental Data Set S3

## Data Availability

Values for seed shape and seed color phenotypes calculated from scans of seeds from each individual plot are provided in Supplemental Data Set S1 with this publication. Python notebooks with code used to generate ground truth data, conduct inference, and calculate model performance are provided at a GitHub repository associated with the study https://github.com/NikeeShrestha/SorghumSeedSegmentation. Cropped seed scans for each image generated as part of the project, including images rejected after QC, are provided as part of the GitHub repository associated with this paper.

## Author Contributions

NS, RVM, and JCS conceived the project. KL designed and conducted the field experiment. HM generated and QCed the genetic marker data. NS, MCT, and JVTR processed images. LLC was responsible for generating ground truth data and validation of image analysis results. NS conducted statistical and genetic analyses, generated figures, and drafted the manuscript. All authors contributed to editing the manuscript.

## Acknowledgments

The authors thank Prince Ngiruwonsanga, Abaigeal Aydt, Isabel Sigmon, and Han Tran for their assistance in collecting the data employed in this study.

## Supplemental Data

The following materials are available in the online version of this article.

- **Supplemental Table S1:** Summary statistics of three seed-shape phenotypes, three seed-color phenotypes, and three seed-color-derived principal components extracted from scans of seeds from individual plots.
- **Supplemental Figure S1:** Geographical distribution of a subset of the Sorghum Diversity Panel (n=328).
- **Supplemental Figure S2:** Photo of the sorghum seeds used as color references during manual sorghum seed phenotyping.
- **Supplemental Figure S3:** Distribution of plot-level and genotype-level phenotype measurements for each seed- related phenotype used in the resampling-based GWAS analysis.
- **Supplemental Figure S4:** Comparison between pre-trained models on rice and wheat seeds.
- **Supplemental Figure S5:** Examples of sorghum genotypes where manual qualitative and automated quantitative color measurements disagree.
- **Supplemental Figure S6:** Interaction between genetic marker (Chr01:72,465,237) linked to Y locus and genetic marker identified on chromosome 3 (Chr03:75,096,302) to affect three color channels derived first principal component.
- **Supplemental Data Set S1:** Plot-level seed-related phenotypes extracted from 1,603 individual plot-level seed scans after removal of the plot level outlier values.
- **Supplemental Data Set S2:** Marker trait associations identified in GWAS conducted automatically measured seed shape phenotypes (A), and seed color phenotypes (B), and manually scored seed color classes in sorghum (C).
- **Supplemental Data Set S3:** Sets of sorghum gene models within mapping intervals for the three marker-trait associations for seed color shown in Figure 5B-D.

---

**Table S1.** Summary statistics of three seed-shape phenotypes, three seed-color phenotypes, and three seed-color-derived principal components extracted from scans of seeds from individual plots.

| phenotype | Mean | Median | SD<sup>a</sup> | SE<sup>b</sup> | Minimum | Maximum |
| --- | --- | --- | --- | --- | --- | --- |
| Seed Area | 909.8 | 908.4 | 193.1 | 4.83 | 447.6 | 1488.6 |
| Seed Length | 48.44 | 48.66 | 5.08 | 0.17 | 34.42 | 62.19 |
| Seed Width | 30.55 | 30.77 | 3.92 | 0.09 | 18.54 | 41 |
| Average Blue Intensity | 100.9 | 105.2 | 24.33 | 0.60 | 57.28 | 171.3 |
| Average Green Intensity | 131.3 | 135.4 | 27.35 | 0.68 | 69.89 | 192.5 |
| Average Red Intensity | 160.4 | 162.3 | 21.31 | 0.53 | 94.5 | 207.9 |
| Principal Component 1 | 0 | 4.66 | 41.67 | 1.04 | -88.9 | 101.9 |
| Principal Component 2 | 0 | 0.24 | 7.07 | 0.17 | -25.2 | 27.21 |
| Principal Component 3 | 0 | 0.09 | 2.89 | 0.07 | -11.7 | 11.18 |

<sup>a</sup> Standard Deviation<sup>b</sup> Standard Error

**Figure S1.**
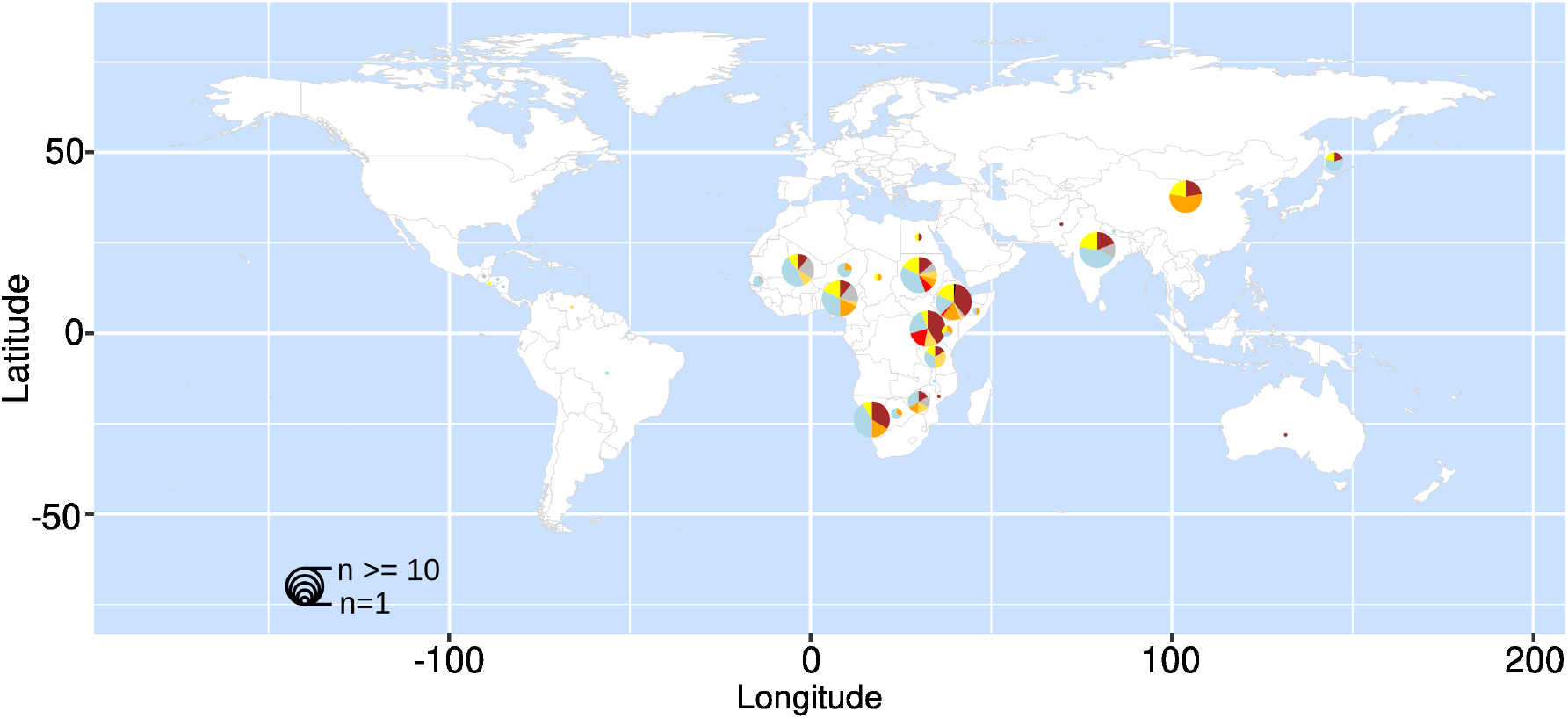
Geographical distribution of a subset of the Sorghum Diversity Panel (n=328). Sorghum genotypes predominantly originated from African countries with diverse seed colors spread across the world. The size of the pie chart varies with the number of genotypes originating from the place where if the number of genotypes *≥* 10, it was assigned the same size. Color mapped to each piechart is based on visual color classification as shown in Figure 2 legend key.

**Figure S2.**
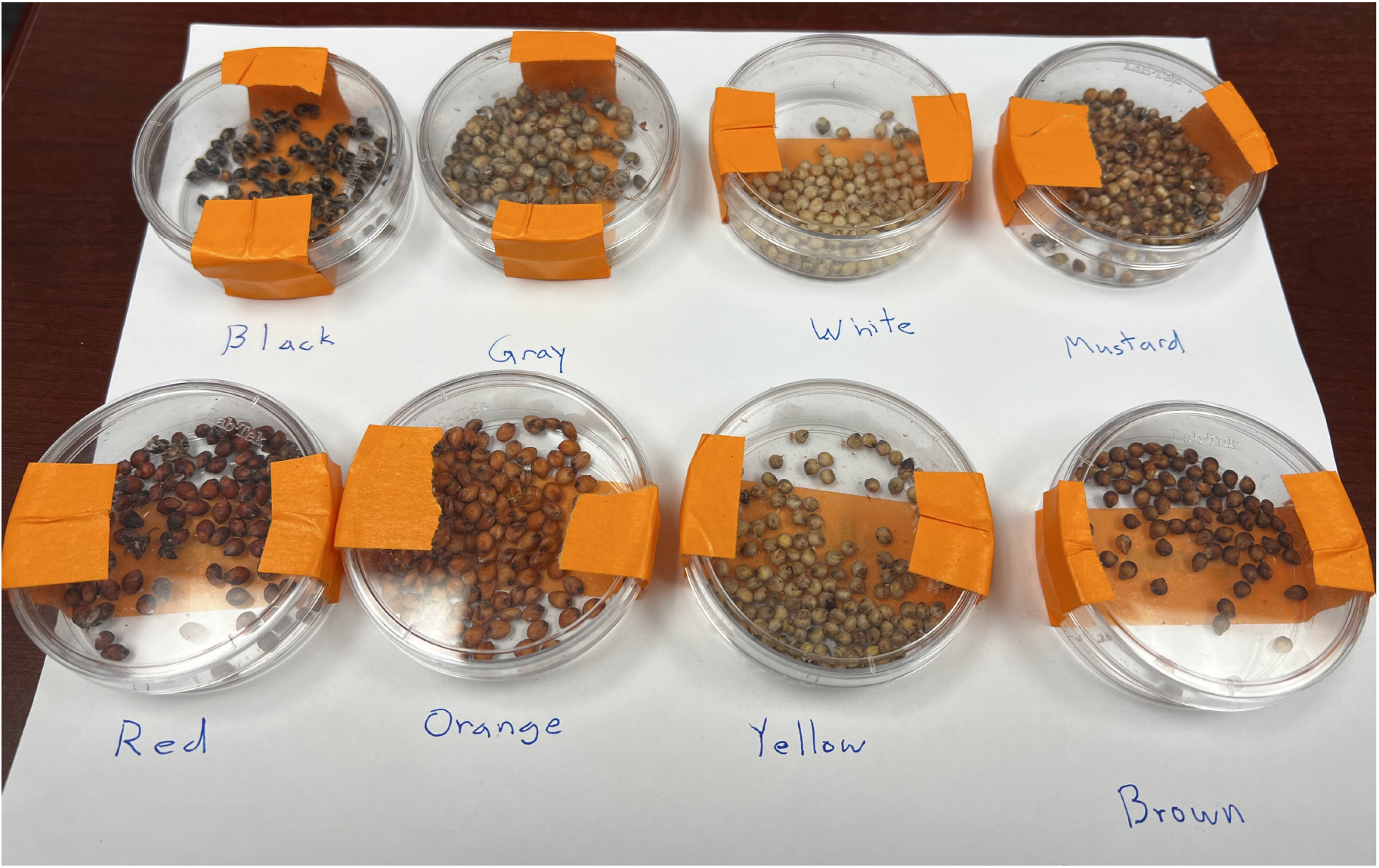
Photo of the sorghum seeds used as color references during manual sorghum seed phenotyping. Seeds were initially scored as belonging to one of eight color classes: white, gray, yellow, mustard, orange, red, brown, and black. These classes were later collapsed into two broader categories: light (white, gray, yellow, mustard) and dark (orange, red, brown, black) for genome-wide association study.

**Figure S3.**
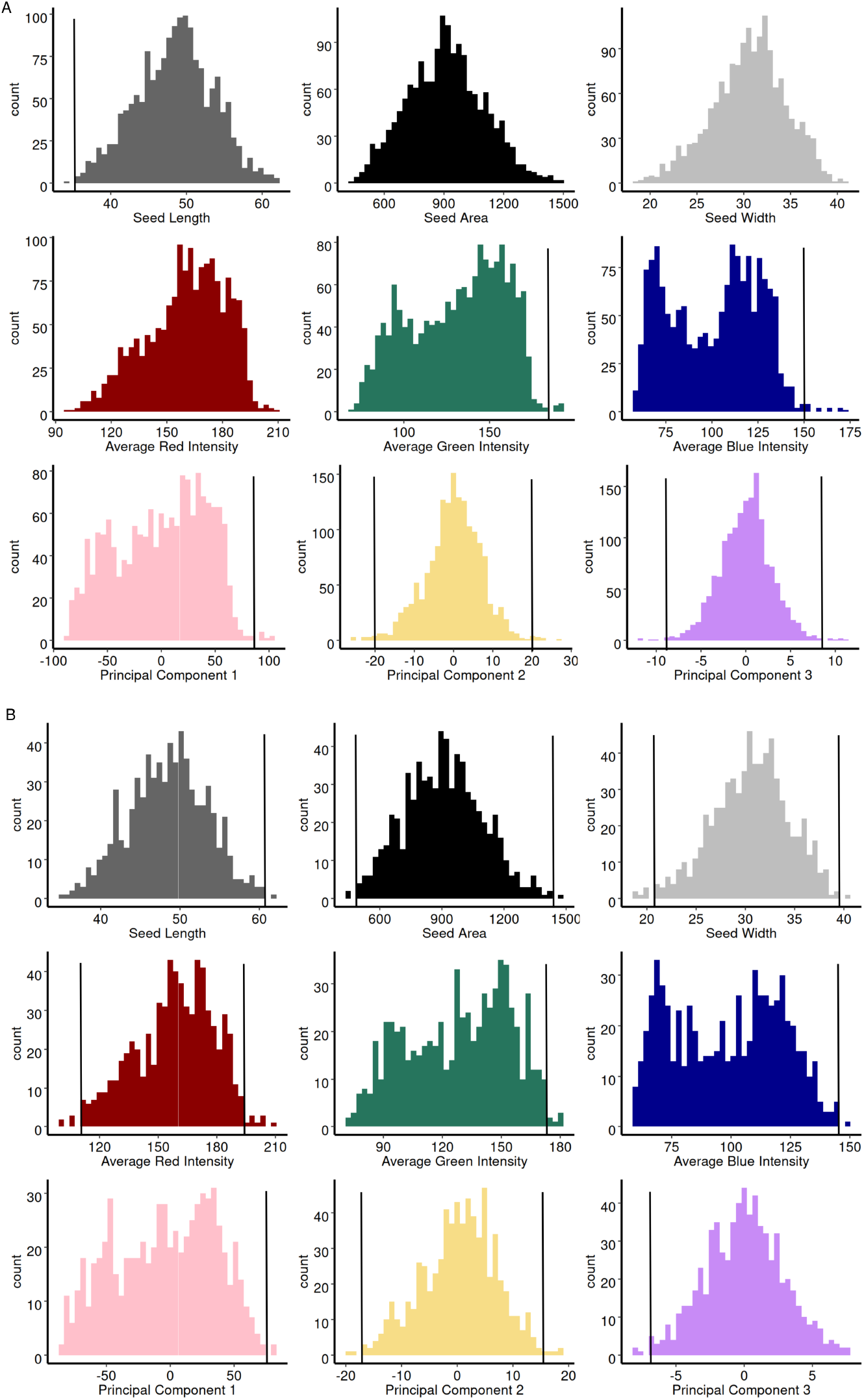
Distribution of plot-level and genotype-level phenotype measurements for each seed-related phenotype used in the resampling-based GWAS analysis. A) Distribution of observed plot level measurements (n = 1,603) for three seed shape phenotypes and six seed color phenotypes. The presence of vertical black lines indicates a cutoff that was applied to a given phenotype to remove extreme values before the calculation of genotype-level values. **B)** Distribution of observed genotype-level average values for 682 sorghum genotypes for which both phenotype and genetic marker data were available. The presence of vertical black lines indicates a cutoff that was applied to a given phenotype to remove extreme values before GWAS analysis.

**Figure S4.**
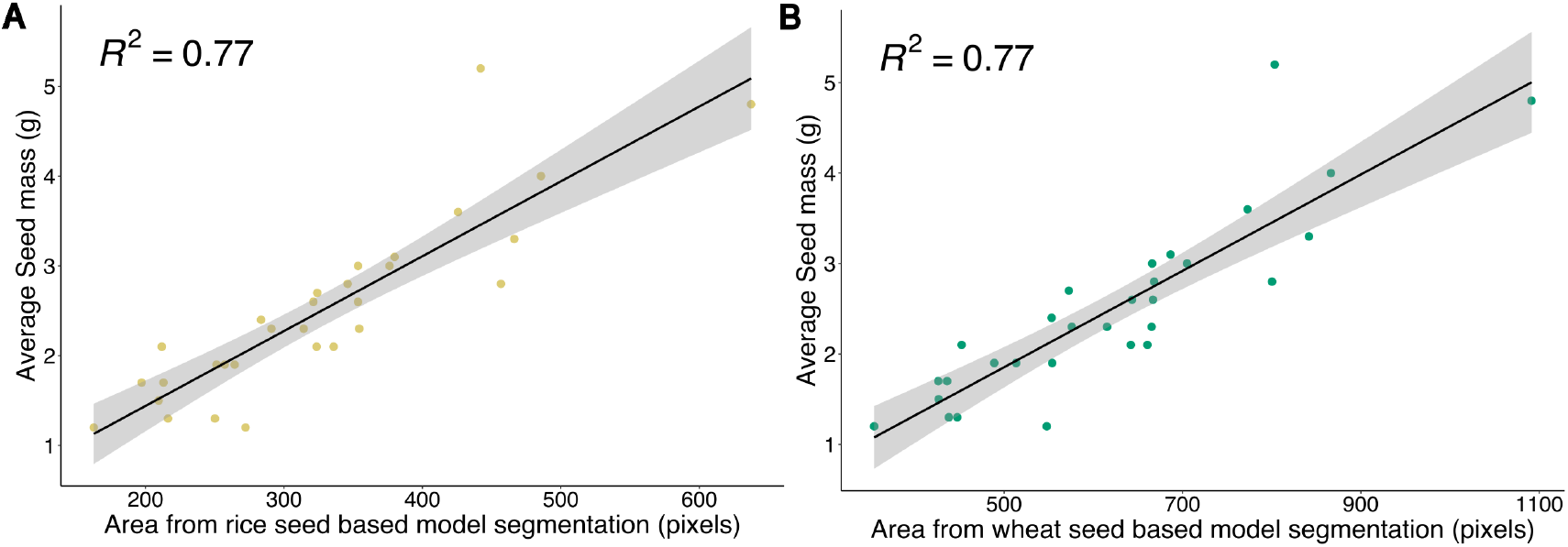
Comparison between pre-trained models on rice and wheat seeds. A) Correlation between area extracted from analyzed new scans of new samples of seed using the rice model (x-axis) and manually measured average seed mass (y-axis) from 30 sorghum genotypes selected to represent the full range of observed seed area distribution observed across scans of all sorghum genotypes included in this study. **B)** Correlation between area extracted from analyzed new scans of new samples of seed using the wheat model (x-axis) and manually measured average seed mass (y-axis) from 30 sorghum genotypes selected to represent the full range of observed seed area distribution observed across scans of all sorghum genotypes included in this study.

**Figure S5.**
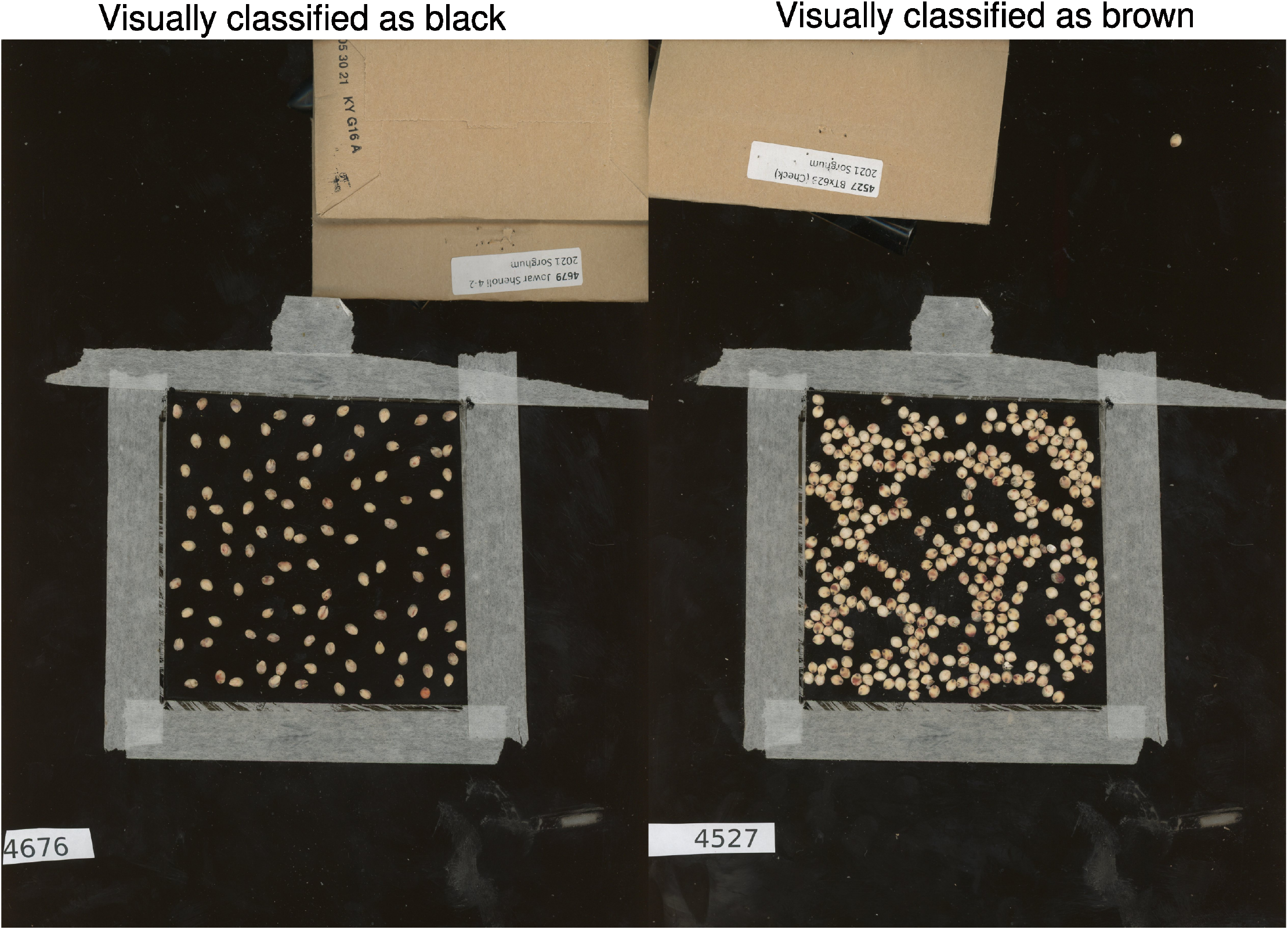
Examples of sorghum genotypes where manual qualitative and automated quantitative color measurements disagree. Sorghum seeds shown above come from two genotypes; SC0499 (left) and BTx623 (right) recorded as "black" and "brown" in manual color classification but not placed in areas of the color space that would correspond to these dark colors based on quantitative color phenotypes (e.g. (Figure 2)).

**Figure S6.**
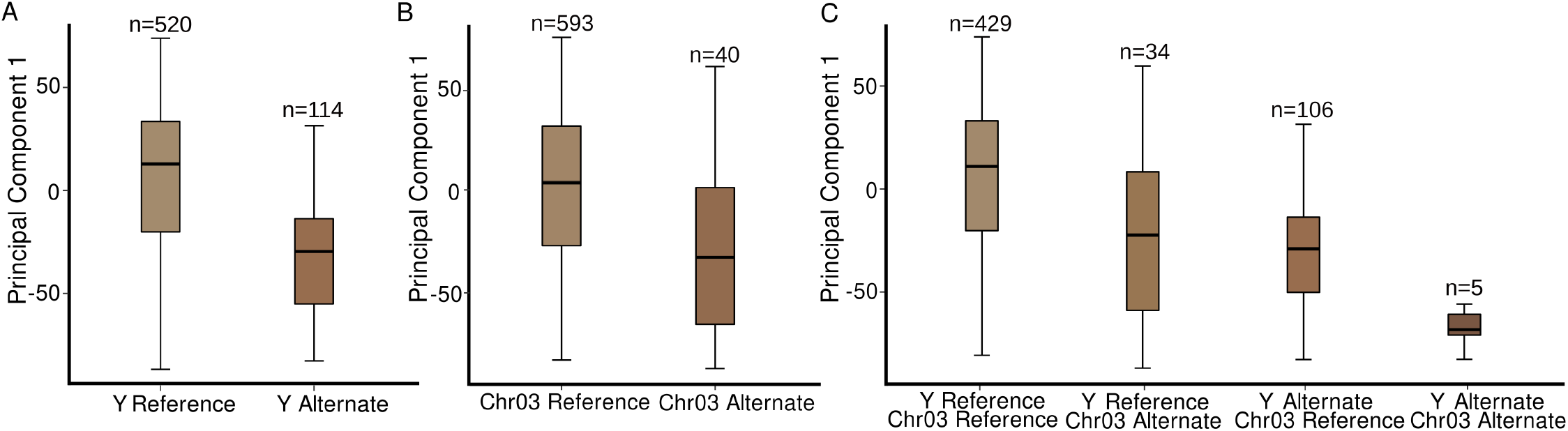
Interaction between genetic marker (Chr01:72,465,237) linked to Y locus and genetic marker identified on chromosome 3 (Chr03:75,096,302) to affect three color channels derived first principal component. A) Difference in scores for the first principal component of variation for sorghum genotypes carrying either the reference or alternative alleles of the most consistently *Y* locus associated GWAS hit (Chr01:72,465,237). For this and subsequent panels sorghum genotypes that carried the alleles of *tan1* and *tan2* adjacent GWAS hits associated with higher tannin concentration were excluded. Box plot colors indicate the median red, green, and blue values for individuals carrying the respective allele. B) Difference in scores for the first principal component of variation for sorghum genotypes carrying either the reference or alternative alleles of the sorghum seed color GWAS hit on chromosome 3.C) Difference in scores for the first principal component of variation for sorghum lines carrying all four possible combinations of homozygous genotypes for the two genetic markers show in panels A and B.

